# The additive polygenic model with assortative mating and shared parent-offspring environment

**DOI:** 10.1101/2022.11.08.515653

**Authors:** Camille Noûs, Hervé Perdry

## Abstract

The consequences of assortative model in the additive polygenic model have been extensively explored by several authors. In this note we extend their results by introducing a correlation between parental and offspring environments.

## 1 Introduction

In this note we assume that the reader is familiar with the additive polygenice model in quantitative genetics, as described in textbooks, e.g [3], [1]. The consequences of assortative mating were considered by Fisher in the very paper in which he proposed the polygenic model [4]. Many other authors have considered the same problem, using different computational approaches, in particular Wright [6], Crow and Felsenstein [2], Nagylaki [5] (see this last paper for a more extensive bibliography).

While assortative mating creates a correlation between mates’ environments, the classical approach consider that the offspring environment is independent from the parental environments. Here we introduce in the model a (positive) correlation between the environmental component of the offspring, and each of its parents environments.

Our results are summarized in the following theorem.

### Theorem 1

*Consider a phenotype determined by P* = *A* + *E, where A is the sum of independent genetic effects, and E is the environmental effect. Consider a couple with two parents indexed by* 1 *and* 2 *and an offspring indexed by* 3. *The values of P, A and E for these individuals are denoted P*_*i*_, *A*_*i*_, *E*_*i*_ *(i* = 1, 2, 3*). Denote* var(*P*) = *σ*^2^, var(*A*) = *a*^2^ *and* var(*E*) = *e*^2^.

*If one assumes that the population is panmictic and that all the environment terms are independent from each other and from the A*_*i*_ ; *then* 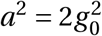 *where* 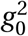 *is the gametic variance, and σ*^2^ = *a*^2^ +*e*^2^ ; *the correlations between parental and offspring genetic effects are* 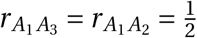.

*We will assume on the contrary that mate choice is partially based on the value of P, so that there is a correlation* 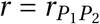 *between the two members of a couple. We also assume that there is a correlation* 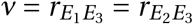 *between the parental and offspring environment, which can be attributed for example to cultural transmission*.

*When r is non-zero, then there is a non-zero gametic correlation r*_*g*_ *between parental gametes; when ν is non-zero, there is a non-zero correlation* 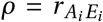 *between the genetic and environmental components. Assuming that the population is at equilibrium, that is, that r*_*g*_ *and ρ are constant accross generations, one can compute r*_*g*_ *and ρ from r, ν*, 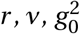 *and e*^2^.

*The presence of parental gametic correlation r*_*g*_ *induces, at equilibrium, a gametic disequilibrium between the causal loci, leading to a gametic variance*

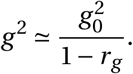

*The above approximation is valid unless r*_*g*_ *is close to* 1, *which in turn occurs only when the trait is almost entirely genetically determined, and when the parental correlation r is close to* 1.

*The additive genetic variance is then* 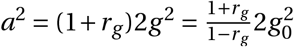. *The correlations between A*_*i*_ *and E* _*j*_ *terms are summarized in the following two tables, which allow to compute the parentoffspring phenotypic correlation*.

The proof of the theorem is the object of the following sections. In section 2 we will derive the consequences of the existence of a gametic correlation *r*_*g*_. In section 3, we consider the components of var(*P*) in an individual in a population in which there is a gametic correlation, as well as a gene-environment correlation. In section 4, we consider a couple with assortative mating and prove the results that are summarized in table 1. In the section 5, we will show how the gametic correlation *r*_*g*_ can be derived from *a, ρ* and *e*, we derive the results summarized in table 2, and we show how, assuming population equilibrium, *ρ* and *ν* are linked. Section 6 gives some technical details about how to compute *ρ* and *a*, given *g*_0_, *e, r* and *ν*.

**Table 1:** Correlations between *A* and *E* terms for the two parents

| | $A_1$ | $E_1$ | $A_2$ | $E_2$ |
| --- | --- | --- | --- | --- |
| $A_1$ | 1 | $\rho$ | $\frac{r}{\sigma^2}(a + \rho e)^2$ | $\frac{r}{\sigma^2}(\rho a + e)(\rho e + a)$ |
| $E_1$ | $\rho$ | 1 | $\frac{r}{\sigma^2}(\rho a + e)(\rho e + a)$ | $\frac{r}{\sigma^2}(\rho a + e)^2$ |
| $A_2$ | $\frac{r}{\sigma^2}(a + \rho e)^2$ | $\frac{r}{\sigma^2}(\rho a + e)(\rho e + a)$ | 1 | $\rho$ |
| $E_2$ | $\frac{r}{\sigma^2}(\rho a + e)(\rho e + a)$ | $\frac{r}{\sigma^2}(\rho a + e)^2$ | $\rho$ | 1 |

**Table 2:** Correlations between *A* and *E* terms for parent / offspring pairs. Here *i* = 1 or 2.

| | $A_3$ | $E_3$ |
| --- | --- | --- |
| $A_i$ | $\frac{1}{2}(1 + \frac{r}{\sigma^2}(a + \rho e)^2)$ | $\rho$ |
| $E_i$ | $\frac{1}{2}(\rho + \frac{r}{\sigma^2}(a + \rho e)(\rho a + e))$ | $v$ |

## 2 Gametes

Assume there are *N* causal loci, with *N* a very large number. The effect of a (haploid) gamete is 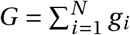, where *g*_*i*_ is the infinitesimal effect of locus *i*. The genetic effect for an individual can be decomposed in *A* = *G*^*p*^ + *G*^*m*^, where *G*^*p*^ (respectively *G*^*m*^) is the effect of the paternal (respectively maternal) gamete.

### 2.1 Inter loci correlation

Let 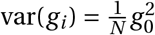and 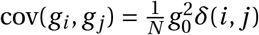(for *i* ≠ *j*). If one has *δ*(*i, j*) = 0 for all *i ≠ j*, then var 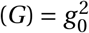. In general,

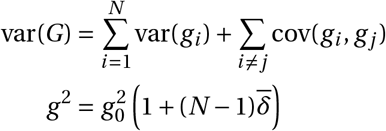

where 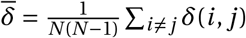. Note that the *δ*(*i, j*) are infinitesimal (they have order of magni-tude 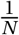).

### 2.2 Méiosis

Consider an individual with genetic component *A* = *G*^*p*^ +*G*^*m*^. One has 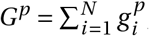, and 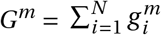. A simple way to model the meiosis is to let *I*_*p*_ ∪ *I*_*m*_ = [1,…, *N*] with 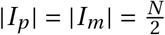. The gamete transmitted has total effect

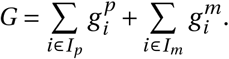

By letting

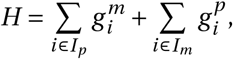

one gets *A* = *G* + *H* ; *H* can be seen as the “untransmitted gamete”.

### 2.3 Evolution of the inter-loci correlations *δ*_*t*_ (*i, j*)

Here we consider the evolution between generation *t* and *t* + 1, assuming that there is a correlation *r*_*g*_ = cor(*G*^*p*^, *G*^*m*^) between the parental gametes.

At generation *t*, individuals are formed by gametes with variance 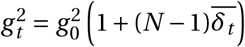.

The two gametes emitted by the two members of a couple in generation *t* − 1 have covari-ance 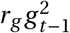. Thus, an individual at generation *t* is formed by two gametes *G*^*p*^ and *G*^*m*^ with covariance 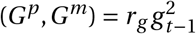. Now 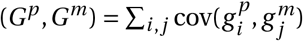, and all the terms in this sum play symmetric roles, hence

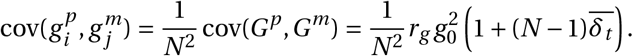

Next, let us consider a gamete emitted by this individual. Let *I*_*p*_ and *I*_*m*_ be as in subsection 2.2, and let 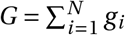 where 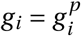 if *i* ∈ *I*_*p*_ and 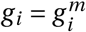 if *i* ∈ *I*_*m*_.

If *i, j* ∈ *I*_*p*_, then 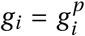 et 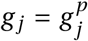 and 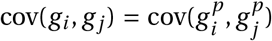, thus *δ*_*t*+1_(*i, j*) = *δ*_*t*_ (*i, j*). The same holds if *i, j* ∈ *I*_*m*_. The remaining cases are *i* ∈ *I*_*p*_, *j* ∈ *I*_*m*_ and *i* ∈ *I*_*m*_, *j* ∈ *I*_*p*_. In both cases, 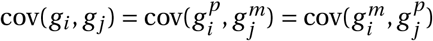, hence

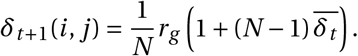

Hence the distribution of *δ*_*t*+1_(*i, j*) is a mixture between *δ*_*t*_ (*i, j*) and this term. We have

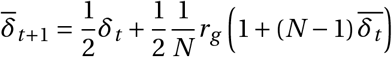

hence at equilibrium, 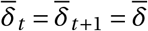 and

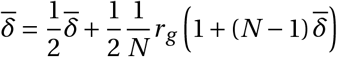

from which one gets

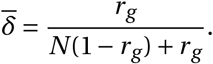

The mean inter-loci correlation is infinitesimal (of order 1/*N*), unless *r*_*g*_ is very close to one. One gets

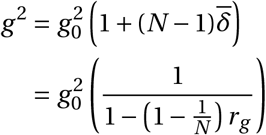

And if 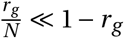, which is the case again unless *r*_*g*_ is very close to one,

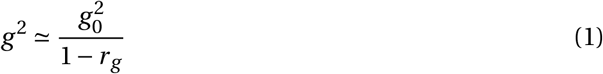

The correlation between parental gametes increases the gametic variance, by inducing an infinitesimal correlation between all the causal loci. Moreover, one has

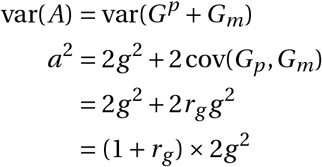

and finally

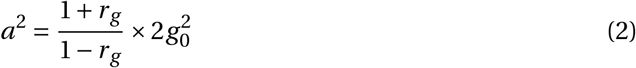

The additive genetic value of an individual is further increased by a factor (1 + *r*_*g*_).

### 2.4 Correlation between transmitted and “untransmitted” gametes

We consider a population at equilibrium. As in subsection 2.2, an individual with additive genetic value *A* = *G*^*p*^ + *G*^*m*^ transmits a gamete 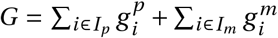, and 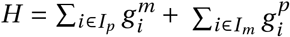 is the untransmitted gamete. Then *A* = *G* + *H* and *G* et *H* play symmetric roles.

The gametic variance is var(*G*) = var(*H*) = *g* ^2^, and 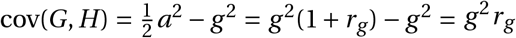, hence the correlation between *G* and *H* is *r*_*GH*_ = *r*_*g*_.

Moreover, one has cov(*A,G*) = cov(*G* + *H, G*) = (1 + *r*_*g*_)*g* ^2^ ; the distribution of *G* conditional to *A* has expected value 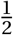*A* and variance 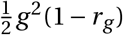.

## 3 A single individual in the population

We consider a population at equilibrium, in which there is a correlation *ρ* = cor(*A, E*) = *r* _*AE*_ between *A* and *E*. We will derive expressions for *r* _*AP*_ and *r*_*EP*_ that will be usefull later.

We have

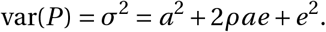

Hence

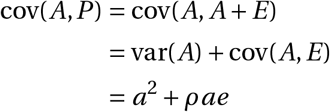

and

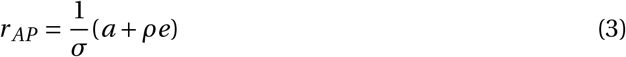

In the same way,

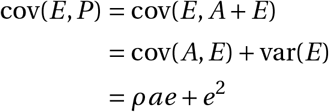

And

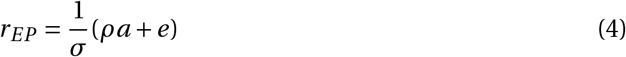

## 4 A couple with assortative mating

In this section and the following, we use the same computational approach than Nagylaki [5]. Consider the two members of a couple, indexed by *i* = 1, 2, with phenotypic correlation 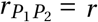. We still have 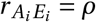 for *i* = 1, 2.

It is assumed that the mate choice is based only on the phenotype. This means that, conditionnal to *P*_2_, *P*_1_ and *A*_2_ are independant, that is 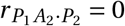 (where 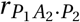 denotes the correlation between *P*_1_ and *A*_2_ conditional to *P*_2_) ; and also 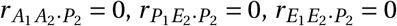.

Then, using lemma 1 (appendix A) and 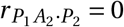, we have

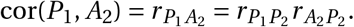

We have (equation 3) 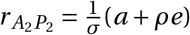, hence

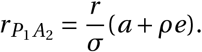

The two members of the couple play symmetric roles so one has 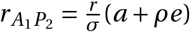.

Applying again lemma 1 to 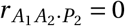, we have

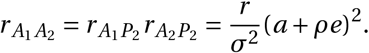

From 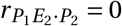 we get

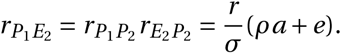

The symmetry argument gives 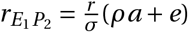.

And from 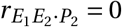, we get

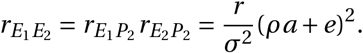

Then we have

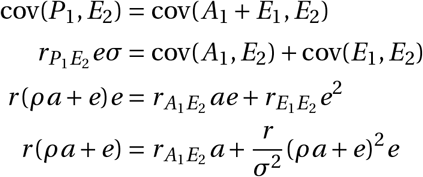

hence

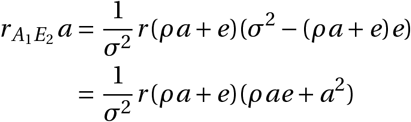

and finally

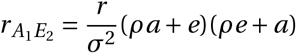

which is also the value of 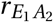.

We have proved all the correlations of table 1.

## 5 Offsprings in assortative mating

### 5.1 Gametic correlation

Assume the individuals *i* = 1, 2 emit gametes *G*_*i*_, with *A*_*i*_ = *G*_*i*_ + *H*_*i*_, as in subsection 2.2. We will link cor 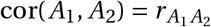 with the gametic correlation *r*_*g*_ = cor(*G*_1_,*G*_2_).

We write

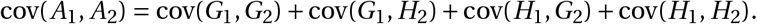

All four terms on the right hand side are equal, and 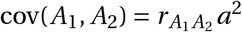, so

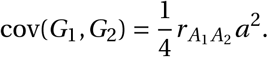

One has *a*^2^ = 2(1 + *r*_*g*_)*g* ^2^ and cov(*G*_1_,*G*_2_) = *r*_*g*_ *g* ^2^, hence 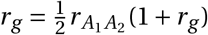 and finally

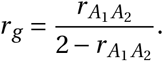

Equivalently,

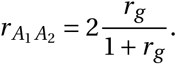

Equation 2 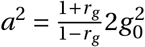 can be written

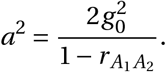

This result is identical to what is derived, for example, in [2]. The approximations made to express *g* ^2^ and, consequently, *a*^2^, were valid unless *r*_*g*_ was very close to one. Here we see that this happens only in the marginal case were 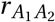 is very close to one, which in turns implies *r* close to one and *e*^2^ ≪ *σ*^2^ : the assortative mating is almost complete, and the environment has almost no part in the trait.

### 5.2 Genetic component

We have *a*^2^ = var(*A*_1_) = cov(*A*_1_,*G*_1_) + cov(*A*_1_, *H*_1_); the two terms on the right hand side are equal, so 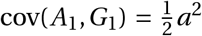. In the same way, from cov(*A*_1_, *A*_2_) = cov(*A*_1_,*G*_2_) + cov(*A*_1_, *H*_2_) we get 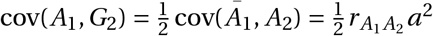.

The additive genetic component of the offspring is *A*_3_ = *G*_1_ + *G*_2_ (one can also write 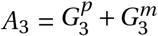 with 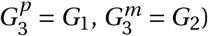. Hence

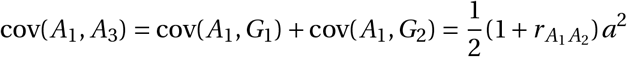

and

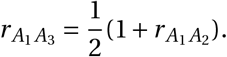

Of course, the two parents play symmetric role and 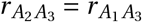.

### 5.3 Environmental component

Let us formalize our assumptions on environmental correlations. We denote 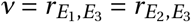. All variables have expected value 0, and var(*E*_*i*_) = *e*^2^ for *i* = 1, 2, 3. The variance-covariance matrix of (*E*_1_, *E*_2_, *E*_3_) is

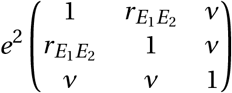

More over, conditionnaly to (*E*_1_, *E*_2_), *E*_3_ is independent from *A*_1_ and *A*_2_.

Note that *ν* can’t be too high; for example if 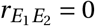, the environment of the offspring can’t be identical to both *E*_1_ and *E*_2_ ; the maximum value in that case is 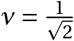. In general, the above matrix is positive definite if

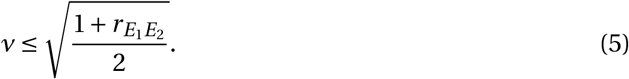

The value 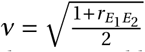 corresponds to the extreme case in which the offspring’s environment is entirely determined by its parents’ environments.

Let us compute cov(*E*_1_, *A*_3_) = cov(*E*_1_,*G*_1_) + cov(*E*_1_,*G*_2_). We have cov(*E*_1_, *A*_1_) = cov(*E*_1_,*G*_1_) + cov(*E*_1_, *H*_1_) ; the two terms in the right hand side are equal, so 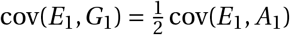. In the same way, 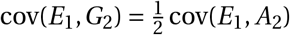. Hence

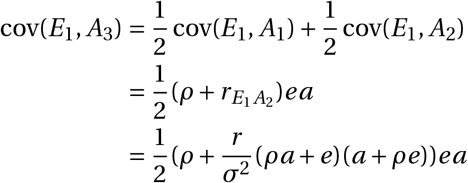

One has 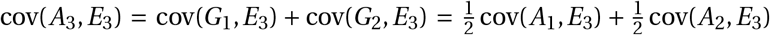. As the two parents play symmetric roles, cov(*A*_1_, *E*_3_) = cov(*A*_2_, *E*_3_) and we get cov(*A*_3_, *E*_3_) = cov(*A*_1_, *E*_3_), hence 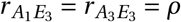, the population beeing at equilibrium.

We have proved all the results summarized in table 2.

### 5.4 Linking *ρ* and *ν*

We are going to compute 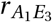 in an other way, linking to *ν*. From the assumption of population equilibrium, we derived 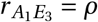; the computation here is the key result allowing to compute the value *ρ* from *ν* and the other parameters of the model.

Conditionnaly to (*E*_1_, *E*_2_), *A*_3_ and *E*_3_ are independent. We are going to apply lemma 2 (appendix A), with *X* = (*A*_1_, *E*_3_) and 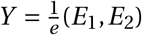. One has

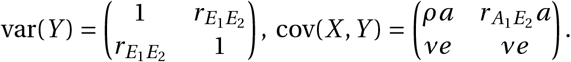

From lemma 2, we get

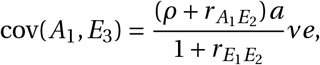

hence

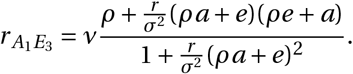

Hence, we have

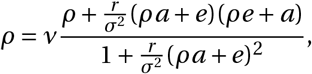

which can be rewritten as

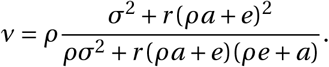

In the next section we show how this relation can be used to compute *ρ*, knowing *ν, g*_0_ and *e*.

## 6 Computing *ρ* and *a*

The natural parameter of the model are *g*_0_ (gametic standard deviation when there is no homogamy), *e* (environmental standard deviation), *r* (correlation between parental phenotypes) and *ν* (environmental correlation between parent and offspring). However we have expressed everything using two extra parameters, *a* and *ρ*. We outline here how these two parameters can be computed from the four aforementioned parameters.

We first define *a* as a function of *ρ* and the other parameters, excepted *ν*. We have 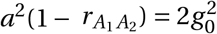 where 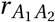 is

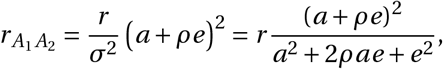

and thus

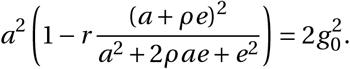

This gives an equation in *a* and *ρ* that can be solved numerically; it has a unique solution 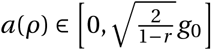. Then, ρ can be computed by solving

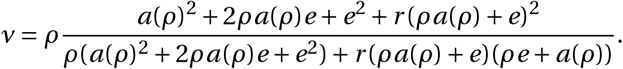

Note that the right hand term varies from 0 for *ρ* = 0 to 1 for *ρ* = 1. Once the equation is solved for *ρ*, one naturally takes *a* = *a*(*ρ*). After this, one needs to verify that the value of *ν* is acceptable, that is if the constraint of equation 5 is satisfied.

As a final remark, note that it is easy to check that if (*g*_0_, *e, r, ν, a, ρ*) satisfy the above constraints, then so does (*κg*_0_, *κe, r, ν, κa, ρ*), for any *κ* > 0, which corresponds merely to a rescaling of the phenotype.

In figure 1 we show *ρ* and *a* as functions of *ν*, for 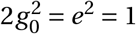 (corresponding to a phenotype with heritability *h*^*2*^ *=* 0.5 when *r = ν =* 0), for several values of *r*. The point at the end of each line corresponds to the maximal value that *ν* can take (see section 5.3).

**Figure 1:**
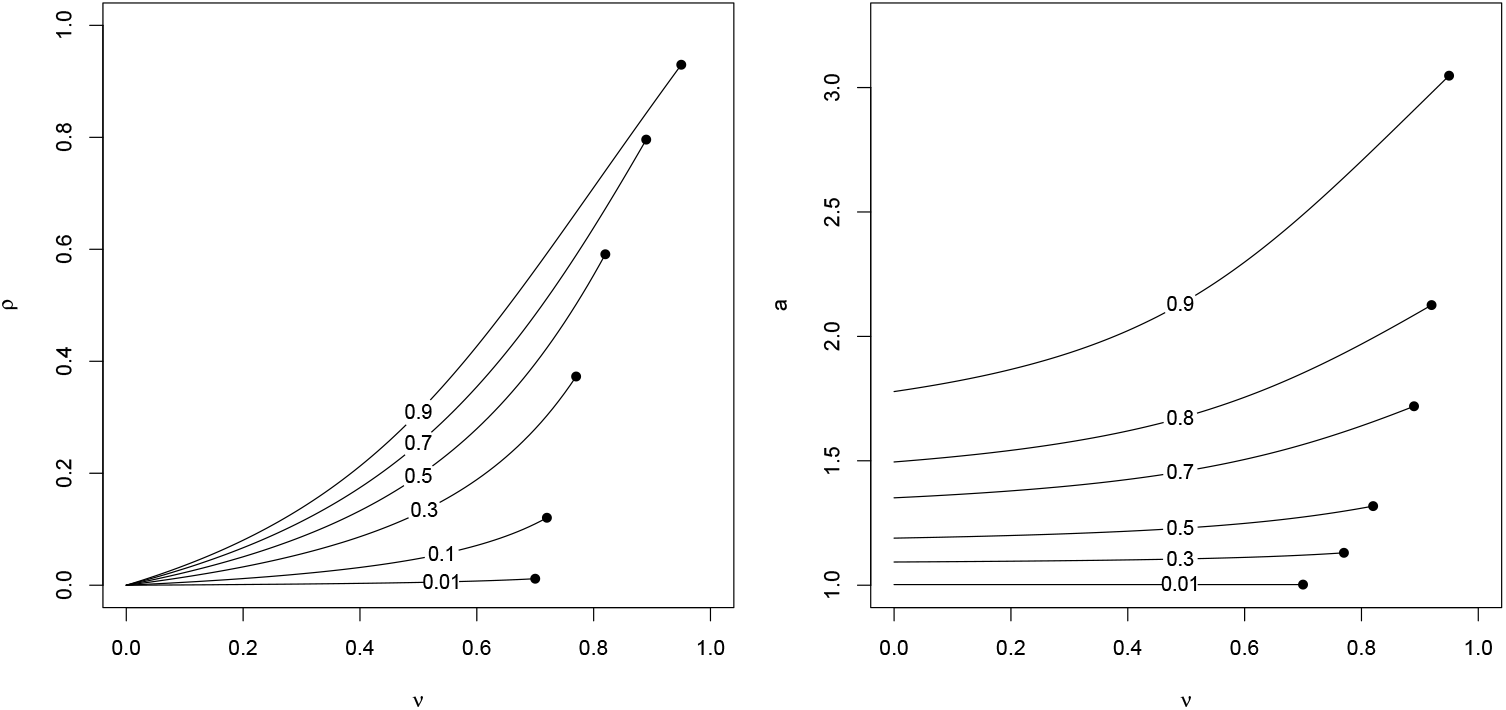
Relation between *ν* and *a, ρ*, for 2*g* 0^2^ = *e*^1^ = 1, and *r* varying from *r* = 0.01 to *r* = 0.9.

## A Technical lemma

If *A, B, X* are three random variables, the partial correlation *r* _*AB* ·*X*_ is the correlation of *A* and *B* conditionnal to *X*.

### Lemma 1

*Let* (*A, B, X*) *be a Gaussian vector. If A and B are independent conditionnal to X, or equivalently r* _*AB* ·*X*_ = 0, *then r* _*AB*_ = *r* _*AX*_ *r*_*BX*_.

**Proof** Assume without lost of generality var(*A*) = var(*B*) = var(*X*) = 1. The variance-covariance matrix of (*A, B, X*) is

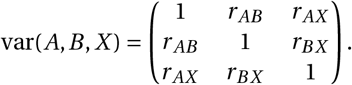

Standard Gaussian theory allows to compute var(*A, B* | *X*):

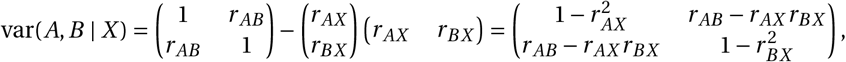

implying *r* _*AB* ·*X*_ = *r* _*AB*_ − *r* _*AX*_ *r*_*BX*_.

### Lemma 2

*Let* (*X*_1_, *X*_2_, *Y*_1_, *Y*_2_) *be a gaussian vector. Note X* = (*X*_1_, *X*_2_) *and Y* = (*Y*_1_, *Y*_2_). *Assume*

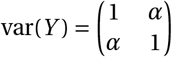

*and*

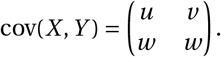

*If X*_1_, *X*_2_ *are independent conditionally to Y, or* cov(*X*_1_, *X*_2_ | *Y*) = 0, *then*

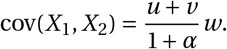

**Proof** The variance-covariance matrix of *X* conditional to *Y* is

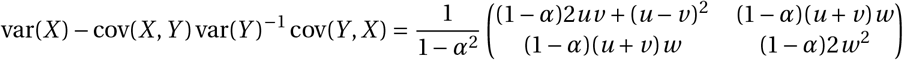

from which the result is obtained readily.

